# On the genetic and environmental reasons why intelligence correlates with criminal victimization

**DOI:** 10.1101/113712

**Authors:** Brian B. Boutwell, Eric J. Connolly, Nicole Barbaro, Todd K. Shackelford, Melissa Petkovsek, Kevin M. Beaver

**Affiliations:** School of Social Work, Saint Louis University; Department of Epidemiology, Saint Louis University; Department of Criminal Justice, Pennsylvania State University, Abington; Department of Psychology, Oakland University; Department of Criminal Justice, University of Central Missouri; College of Criminology and Criminal Justice, Florida State University

**Author notes:** Please address correspondence to: Brian B. Boutwell, Ph.D. School of Social Work College for Public Health & Social Justice Saint Louis University 3550 Lindell Boulevard St. Louis, MO 63103-1021, USA.

**Keywords:** intelligence, victimization, criminology, behavioral genetics

## Abstract

Researchers have expended considerable effort to understand the causes and correlates of criminal victimization. More recently, scholars have focused on identifying individual-level traits that increase the odds of victimization. Generally absent from this line of research, however, is examining the extent to which previously unmeasured genetic and environmental influences contribute to the covariation between victimization and individual-level risk factors. The current study aims to replicate and extend prior research by examining the contribution of genetic and environmental influences on the association between intelligence and victimization by analyzing twin and sibling data from two nationally representative samples of American youth. Quantitative genetic analyses indicate that common additive genetic factors, as well as non-shared environmental factors, explained the phenotypic association between intelligence and victimization. Finally, our results revealed that after correcting for possible familial confounding, the effect of intelligence on victimization experiences remained statistically significant. The findings of the current study replicate and extend prior research on the phenotypic association between indicators of general intelligence and the experience of victimization.

---

Personal victimization can have serious consequences, ranging from physical injury and loss of property, to psychological and emotional trauma (Graham & Juvonen, 2001; Menard, 2002). Researchers across a range of disciplines, including criminology, psychology, and sociology, have expended considerable effort to understand the factors that lead to personal victimization, and to construct theories of victimization that integrate findings from this body of research (Cohen and Felson, 1979). To date, however, much of the effort devoted to understanding the risk factors for victimization has focused on identifying *environmental* factors, such as lifestyle choices, residence in disadvantaged neighborhoods, and exposure to delinquent peers (routine activities/lifestyle theories; e.g., Averdijk, 2011; Schreck & Fisher, 2004; Schreck, Fisher, & Miller, 2004) to explain personal victimization.

Recently, however, studies have begun examining individual-level attributes that may increase the likelihood of personal victimization. Certain cognitive factors and psychological traits, for instance, have been found to increase the risk of being criminally victimized ( Beaver et al.,2016; Cohen & Felson, 1979). Self-control, for example, is consistently associated with an increase in the odds of victimization (see Pratt, Turanovic, Fox, & Wright, 2014, for meta-analysis). Several other psychological traits, such as anger, psychopathy, and self-regulation (Finkelhor, Ormrod, & Turner, 2007; Henry, Caspi, Moffitt, & Silva, 1996; Silver, Piquero, Jennings, Piquero, & Leiber, 2011) have also been identified as increasing the risk of personal victimization.

Given the consistent association between victimization and traits such as self-control, scholars have also hypothesized that another individual-level trait—general intelligence—may predict victimization. This hypothesis is derived from research findings indicating that lower intelligence is associated with exposure to criminal peers (Kimonis, Frick, & Barry, 2004; Seals & Stern, 2013), drug culture (Duncan, Kennedy, & Smith, 2000; Latvala et al., 2009), lower self-control (Meldrum et al., 2017), and greater risk-taking propensities (Pharo, Sim, Graham, Gross, & Hayne, 2011)—all of which are associated with increased odds victimization. Beaver and colleagues (2016) directly tested the intelligence-victimization link using nationally representative data from the National Longitudinal Study of Adolescent to Adult Health and documented an association between verbal intelligence and criminal victimization in early adulthood. The results suggest that as scores on an indicator of general intelligence decreased, the odds of falling victim to criminal behavior increased.

Additionally, a growing line of research has begun examining the potential for genetic contributions to victimization experiences (Vaillancourt, Hymel, & McDougall, 2013) by utilizing behavioral genetic designs—adoption, twin, or sibling analyses. These studies have consistently revealed that victimization experiences are at least moderately heritable (Ball et al., 2008; Connolly & Beaver, 2014; Barnes & Beaver, 2012; Beaver, Boutwell, Barnes, & Cooper, 2009; Beaver et al., 2007; Hines & Saudino, 2004). This is particularly important given that the phenotypes correlate with victimization are also, to varying degrees, heritable (see Beaver et al., 2016). Individual differences in levels of selfcontrol, for example, has consistently been shown to be under genetic influence (Beaver, Wright, DeLisi, & Vaughn, 2008). Criminal involvement is also moderately to highly heritable (Barnes, Beaver, & Boutwell, 2011). General intelligence, likewise, is highly heritable—becoming increasingly so as individuals age (Plomin & Deary, 2015). Taken together, this evidence suggests that at least part of the reason why victimization covaries with other phenotypes could be because shared genetic factors influence both traits (see Barnes et al., 2014 for more discussion).

Despite the possible genetic correlation between victimization and associated risk factors, little empirical work has directly addressed the issue of shared genetic etiology for explaining victimization, more generally. Barnes and Beaver (2012) examined the victim-offender overlap using a nationally representative sample and reported that genetic factors explained between 51% and 98% of the association between offending and victimization. Similar results have been documented for the association between victimization and low self-control (Boutwell et al., 2013), and for the association between violent victimization and criminal behavior (Vaske, Boisvert, & Wright, 2012). It is therefore reasonable to hypothesize that the effect of certain risk factors, such as intelligence, on victimization might exist not because one is necessarily causing the other, but also because the two traits correlate at a genetic level. Both general intelligence and victimization are heritable traits and, as mentioned above, Beaver and colleagues (2016) documented a negative association between verbal intelligence and criminal victimization experiences in early adulthood. What the analysis by Beaver and colleagues (2016) did not reveal was the extent to which the phenotypic correlation between intelligence and victimization may be accounted for by shared—or correlated—genetic and environmental influences.

The current study aims to replicate and extend prior research on the common genetic and environmental influences on intelligence and victimization. We analyzed data from two nationally representative samples of American youth to first estimate the phenotypic correlation between indicators of general intelligence and criminal victimization. If both traits correlate at the phenotypic level—which we hypothesize based on findings from Beaver et al. (2016)—we will then extend the analysis and estimate a series of behavioral genetic models to examine the relative contribution of genetic and environmental influences on the phenotypic correlation between multiple indicators of general intelligence and criminal victimization.

## Method

### Data

To further examine the link between indicators of general intelligence and victimization, we utilized data drawn from the National Longitudinal Survey of Youth 1997 (NLSY97) and the Children of the National Longitudinal Survey of Youth 1979 (CNLSY).

#### NLSY97 Sample

The NLSY97 is a nationally representative sample of youth born between 1980 and 1984 living in the United States during the initial survey wave (see also, Connolly & Beaver, 2015; 2016). Participants were first surveyed in 1997, and then assessed annually from 1997 to 2011. The NLSY97 sample was the product of a stratified multistage cluster probability sampling design where over 90,000 households were initially selected using probability sampling methods. After this step, NLSY staff identified a target sample of 9,808 age-eligible youth for participation in the study. Youth between the ages of 12 and 16 years as of December 31, 1996 were asked to participate in the NLSY97. There were multiple youth between the ages of 12 and 16 years from the same household who agreed to participate in the NLSY97, resulting in many participants in the NLSY97 being biologically related to one another. Previous research has taken advantage of questions asking respondents about their biological or social relationship with other household members to identify levels of biological relatedness between participants in the NLSY97 (Connolly & Beaver, 2016). To validate the kinship links from this method, a series of biometric analyses were conducted using measures of height. Because height is a highly heritable phenotypic trait (heritability estimates ranging from *h^2^* = .80 to *h^2^* = .90; Silventoinen et al., 2003), height scores were standardized by average heights in the NLSY97 based on age and sex sample norms. Results indicated strong convergent validity between heritable estimates for male and female height in the NLSY97 and those reported in other heterogeneous sibling samples (for more information, see Connolly & Beaver, 2016).

Once sibling pairs of varying levels of genetic relatedness in the NLSY97 were identified, one sibling pair per household was randomly selected to be included in the sample. Because the NLSY97 is a nationally representative sample of youth and staff did not over sample for twins, full siblings represent close to 90% of the sibling sample. A random sample of full-sibling pairs was therefore taken from this population to be included in the final NLSY97 sibling sample. After randomly selecting a sample of full-sibling pairs, the final analytic sample included *n* = 1,085 sibling pairs that included 22 monozygotic (MZ) twin pairs (who share 100% of their segregating genes), 30 dizygotic (DZ) twin pairs (who share, on average, 50% of their segregating genes), 947 full-sibling pairs (who share, on average, 50% of their segregating genes), and 86 half-sibling pairs (who share, on average, 25% of their segregating genes). All siblings included in the final analytic sample provided valid information on each measure examined in the current study.

#### CNLSY Sample

The CNLSY is a sample of youth born to a nationally representative sample of women between the ages of 14 and 21 years in 1979 (NLSY79). Beginning in 1986, children born to women from the NLSY79 were surveyed to create the CNLSY. Children in the CNLSY have been surveyed biennially, beginning in 1986, and completed measures designed to assess cognitive, emotional, and behavioral development. In 1994, children age 15 years and older were administered a self-report survey that asked questions about age-appropriate behaviors, including sexual intercourse, deviance, personal relationships, substance use, and victimization. Because the CNLSY sampled multiple children from the same mother, many participants are biologically related to one another. Although information about levels of biological relatedness between participants was not originally collected, Rodgers et al. (1994) has used self-report information on the type of relationship shared between each participant and other household members to develop a linking algorithm that assigns children from the CNLSY a sibling status and genetic relatedness score. The kinship links developed by Rodgers and colleagues (1994) have been validated by a series of tests, and have been used as reliable indicators of genetic relatedness for CNLSY children in over 45 peer-reviewed publications (Rodgers et al., 2016). Using these kinship links, a sibling sample of *n* = 2,854 sibling pairs including 1,532 full-sibling pairs and 1,322 half-sibling pairs with information on all key variables of the current research were identified.

### Measures

#### Intelligence

Intelligence for siblings from the NLSY97 was assessed using scores from the computer-adaptive form of the Armed Services Vocational Aptitude Battery (ASVAB). The ASVAB was administered from the summer of 1997 through the spring of 1998 when siblings in the analytic sample were between the ages of 12 and 18 years. The ASVAB measures the participant’s knowledge and skills in a variety of topical areas including arithmetic reasoning, mathematics knowledge, paragraph comprehension, and word knowledge. The ASVAB is a well-validated and reliable measure of intelligence and has been found to predict academic achievement and job performance (Palmer, Hartke, Ree, Welsh, & Valentine, 1988; Welsh, Kucinkas, & Curran, 1990). NLSY staff created sample weights based on respondent age and assigned percentiles for the theta scores for tests assessing knowledge for the noted topical areas. ASVAB scores were then transformed into percentile scores based on an aggregated, intra-group normed percentile score and sample weights. Participant scores on the ASVAB ranged from 0 to 99.

Intelligence for siblings from the CNLSY was assessed using scores from three separate sub-tests of the Peabody Individual Achievement Test (PIAT; Dunn & Markwardt, 1970) and scores from the Peabody Picture Vocabulary Test – Revised (PPVT-R; Dunn & Dunn, 1981). The PIAT has been employed widely as an assessment of intelligence in prior research (Baker, Keck, Mott, & Quinlan, 1993; Connolly & Beaver, 2015). The PIAT was administered to subjects in the study when they were between the ages of 5 and 14 years. Reading comprehension in the PIAT was measured via 64 items. Overall, the items assessed how often respondents selected one out of four pictures that accurately explained the meaning of a sentence. Reading recognition, comprised of 84 individual items, measured how often participants correctly recognized printed letters and read words aloud. Mathematics was measured by 84 questions assessing each participant’s knowledge of mathematical concepts applicable for their age (see Connolly & Beaver, 2015 for more detail).

The current research used PIAT scores collected when siblings were between the ages of 13 and 14 years. PIAT scores were standardized to have a mean of 100 with a standard deviation of 15. The PPVT-R was also administered to all CNLSY participants between the ages of 4 and 14 years to measure hearing and receptive vocabulary for Standard American English. Hearing and receptive vocabulary was assessed by NLS interviewers saying a word to participants and asking them to point to 1 of 4 pictures that best portrayed the word’s meaning. The current study used PPVT-R scores that were collected when siblings were between the ages of 13 and 14 years. PPVT-R scores were standardized to have a mean of 100 with a standard deviation of 15.

#### Victimization

During the 2002, 2007, and 2013 NLSY97 survey waves, participants were asked about stressful life events, including the following question: “In the past 5 years, have you been the victim of a violent crime, for example, physical or sexual assault, robbery, or arson?” Participants who responded “yes” to this question during any of the survey waves were given a value of “1,” whereas participants who responded “no” to this question at all survey waves were given a value of “0”. During the 2004, 2006, 2008, 2010, 2012, and 2014 CNLSY survey waves, participants were asked the following question: “Since [the date of the last interview], have you been the victim of a violent crime, for example, physical or sexual assault, robbery, or arson?” Participants who responded “yes” to this question during any of the survey waves were given a value of “1,” whereas participants who responded “no” to this question at all survey waves were given a value of “0”.

#### Control Variables

Four control variables are included in the multivariate regression analyses: Family income, participant age, race, and sex. Family income was measured as a quartile measure that divided siblings’ family net household income (measured in US dollars) into categories based on their ranking in the sample distribution (1 = 1% to 25%, 2 = 26% to 50%, 3 = 51% to 75%, 4 = 76% to 99%). Age was measured as a continuous variable in years. Race was measured with a dichotomous dummy variable (0 = *Non-Black/Non-Hispanic*, 1 = *Black/Hispanic/Mixed Race*). Sex was measured as a dichotomous dummy variable (0 = *female*, 1 = *male*).

### Plan of Analysis

The analyses are conducted in a series of interconnected steps. We first examine the associations between intelligence measures and the probability of victimization in each sibling sample (i.e., NLSY97 and CNLSY) by estimating a series of multivariate binary logistic regression models. All models include measures of family income, age, race, and sex to control for possible confounding variables. To examine whether there are differences in risk for victimization across the distribution of intelligence, we follow Beaver et al. (2016) and estimate the odds of victimization across different quartiles of intelligence. We first estimate a series of binary logistic regression models that examine the risk for victimization using each percentile or standardized measure of intelligence to test whether intelligence scores predict risk for victimization after controlling for family income, age, race, and sex. The second set of binary logistic regression models examines the association between intelligence scores in the bottom 25^th^ percentile (i.e., the lowest quartile of intelligence) and risk for victimization (0 = scores between 26^th^ and 99^th^ percentile, 1 = scores between 1^st^ and 25^th^ percentile). Each model thereafter examines the risk for victimization across different intelligence scores falling in the 50^th^ (second lowest quartile), 75^th^ (second-highest quartile), and top 25^th^ percentile (highest quartile). In accord with previous research (Beaver et al., 2016), we hypothesize that respondents in the lower quartiles of intelligence will be at the greatest risk of victimization, whereas respondents in the top quartiles will have the lowest risk for victimization. Models were estimated using robust standard errors because sibling data were nested within households.

Next, between-sibling correlations for intelligence and victimization are calculated in each sibling sample to examine the extent to which one sibling’s score on intelligence measures or risk for victimization correlates with their co-sibling’s score on the same measure. If between-sibling correlations are larger for MZ twins (who share 100% of their genes) compared to DZ twins and full-siblings (who share, on average, 50% of their genes), and between-sibling correlations are larger for DZ twins and full-siblings compared to half-siblings (who share, on average, 25% of their genes), this can be interpreted as evidence for genetic influence on the measures under examination. Because measures of intelligence and victimization in both sibling samples are dichotomous (0, 1), tetrachoric correlations are calculated. Between-sibling correlations were estimated with standard errors adjusted for nonindependence since siblings from the same family contributed to nested observations between indicators of intelligence and victimization (Asparouhov & Muthén, 2006).

While between-sibling correlations are useful for providing initial insight into whether there is genetic influence on a variable of interest do not provide *precise* heritability and environmental estimates for the variables of interest. The third step in our analysis, therefore, involves estimating a series of univariate ACE liability-threshold models using *Mplus* version 7.1 (Muthén & Muthén, 1998-2011). ACE liability-threshold models provide accurate estimates of the proportion of variance in liability for a variable (e.g., quartile of intelligence or victimization) accounted for by additive genetic influences (symbolized as A), shared environmental influences (symbolized as C), and nonshared environmental influences (including error) (symbolized as E) (Plomin et al., 2013; Prescott, 2004). To generate estimates for A, the correlation between the latent A components for sibling pairs are fixed to accord with levels of genetic relatedness shared between siblings. The correlation between latent A components for MZ twins, therefore, is fixed to 1.0, whereas the correlation between latent A components for DZ twins and full-siblings is fixed to .5, and fixed to .25 for half-siblings. The correlation between latent C components was fixed to 1.0 because siblings are assumed to share 100% of the shared environment, and the correlation between latent E components was fixed to 0 because siblings are assumed to share 0% of the nonshared environment. Parameter estimates for the amount of variance in liability accounted for by A, C, and E is computed by comparing observed between-sibling correlations to predicted between-sibling correlations generated by the model.

The next step in the analysis involves estimating a series of bivariate liability-threshold models. These models are estimated to examine the proportion of the covariance between intelligence and victimization (i.e., the extent to which the correlation between intelligence and victimization is accounted for by common genetic influences underpinning both traits) for sibling pairs from the NLSY97 and CNLSY. Estimates for the A, C, and E parameters from the bivariate models reveal the proportion of covariance between two latent variables that is accounted for by additive genetic influences (A), shared environmental influences (C), and nonshared environmental influences (E). Bivariate models are also estimated using *Mplus* version 7.1 (Muthén & Muthén, 1998-2012). All univariate and bivariate ACE liability-threshold models are estimated using the weighted least squares robust estimator option. Model fit is evaluated using an adjusted *χ^2^* difference test (Santorra, 2000) and values from two model fit indices: the comparative fit index (CFI) and the root mean square error of approximation (RMSEA). Acceptable model fit cutoff points for the CFI were values greater than or equal to .90, which indicate satisfactory fit, and values for the RMSEA less than or equal to .05, which indicate good fit (Hu & Bentler, 1999).

The last step in the analysis involved estimating within-sibling logistic regression models to examine the effect of intelligence (measured continuously) on risk for victimization after controlling for influences that make siblings similar to one another and cluster within families, such as genetic and shared environmental factors. Within-sibling regressions assess whether one sibling with a higher or lower score on a measure of interest is significantly more or less likely to report an outcome. In this case, within-sibling regressions were estimated to examine whether siblings with a higher intelligence score were less likely to report being the victim of a violent crime. If siblings with a higher intelligence score were significantly less likely to report being victimized, and the magnitude of this effect did not vary across sibling pairs who share different levels of genetic material, this could be interpreted as evidence that intelligence affects risk for victimization (McGue et al. 2010).

## Results

The first step of the analysis is estimating a series of multivariate binary logistic regression models for victimization.^1^ Estimates from the first model examining the independent effect of ASVAB scores on victimization for the NLSY97 sample are presented on the left-hand side of Table 2. Participants with below average ASVAB scores were more likely to report having been the victim of a violent crime. This effect was independent of family income, age, race, and sex. Model 2 examines the association between respondents with an ASVAB score in the bottom 25^th^ percentile of the distribution and victimization. The results indicate that respondents with an ASVAB score in the bottom 25^th^ percentile were not more likely to be victimized. Model 3, however, indicates that respondents with an ASVAB score in the 50^th^ percentile of the distribution had an increased risk of being victimized. Model 4 reveals that the odds of victimization declined for respondents with an ASVAB score in the 75^th^ percentile and the top 25^th^ percentile.

**Table 1.** Descriptive Statistics

|  | Mean | *SD* | Min | Max |
| --- | --- | --- | --- | --- |
| <u>NLSY97</u> |  |  |  |  |
| ASVAB | 44.74 | 29.81 | 1.00 | 99.00 |
| Victimization | .11 | .17 | 0 | 1 |
| Family Income | 42,352.72* | 35,621.10* | 0.00* | 246,474.00* |
| Age | 31.52 | 1.59 | 29 | 33 |
| Race | .21 | .40 | 0 | 1 |
| Sex | .51 | .46 | 0 | 1 |
| <u>CNLSY</u> |  |  |  |  |
| PIAT Reading Comprehension | 98.04* | 14.05* | 65* | 135* |
| PIAT Reading Recognition | 100.99* | 15.43* | 65* | 135* |
| PIAT Mathematics | 98.76* | 14.21* | 65* | 135* |
| PPVT-R | 97.64* | 16.43* | 20* | 160* |
| Victimization | .08 | .19 | 0 | 1 |
| Family Income | 39,541.56* | 43,836.43* | 0.00* | 164,421.68* |
| Age | 23.76 | 10.76 | 16 | 32 |
| Race | .17 | .36 | 0 | 1 |
| Sex | .52 | .42 | 0 | 1 |

Notes: \* indicates values before transformation

**Table 2.** Results from Binary Logistic Regression Models Predicting Victimization in the NLSY97

| Variable | *Model 1* |  |  | *Model 2* |  |  | *Model 3* |  |  | *Model 4* |  |  | *Model 5* |  |  |
| --- | --- | --- | --- | --- | --- | --- | --- | --- | --- | --- | --- | --- | --- | --- | --- |
|  | *OR* | *SE* | *95% CI* | *OR* | *SE* | *95% CI* | *OR* | *SE* | *95% CI* | *OR* | *SE* | *95% CI* | *OR* | *SE* | *95% CI* |
| **ASVAB** |  |  |  |  |  |  |  |  |  |  |  |  |  |  |  |
| Percentile Measure | .98** | .02 | [.93-.99] | - | - | - | - | - | - | - | - | - | - | - | - |
| Bottom 25th Percentile | - | - | - | 1.01 | .02 | [.96-1.73] | - | - | - | - | - | - | - | - | - |
| 50th Percentile | - | - | - | - | - | - | 1.14** | .03 | [1.02-1.96] | - | - | - | - | - | - |
| 75th Percentile | - | - | - | - | - | - | - | - | - | .93** | .03 | [.65-.99] | - | - | - |
| Top 25th Percentile | - | - | - | - | - | - | - | - | - | - | - | - | .75** | .02 | [.70-.97] |
| Family Income | .92* | .05 | [.83-.98] | .90* | .04 | [.80-.97] | .91* | .04 | [.81-.98] | .91* | .04 | [.81-.98] | .91* | .04 | [.79-.99] |
| Age | 1.01 | .02 | [.96-1.05] | 1.00 | .02 | [.96-1.04] | 1.01 | .02 | [.92-1.09] | .99 | .01 | [.97-1.02] | 1.01 | .02 | [.96-1.07] |
| Race | .91 | .07 | [.84-1.03] | .89 | .06 | [.83-1.02] | .90 | .06 | [.84-1.03] | .90 | .05 | [.83-1.02] | .90 | .06 | [.80-1.05] |
| Sex | .90* | .06 | [.83-.98] | .92* | .04 | [.87-.98] | .92* | .04 | [.85-.98] | .92* | .04 | [.85-.99] | .92* | .05 | [.84-.99] |

Notes: *CI* = confidence interval. \*\* $p < .01$ ; \* $p < .05$

Table 3 presents the results from the second set of binary logistic regression models that examine the association between different quartiles of intelligence and risk for criminal victimization among siblings from the CNLSY. There was a negative and significant association between all standardized measures of intelligence and victimization. On average, siblings with lower reading comprehension, reading recognition, mathematics, and hearing and receptive vocabulary were more likely to report having been the victim of a violent crime. Models 2 through 5 examine the association between different quartiles of intelligence and victimization. The results indicate that the risk for criminal victimization gradually decreased as intelligence increased. Specifically, siblings in the bottom 25^th^ percentile of all measures of intelligence demonstrated the highest risk for victimization, whereas siblings in the top 25^th^ percentile of all measures of intelligence demonstrated the lowest risk for victimization. All effects were independent of family income, age, race, and sex. The results from each multivariate logistic regression model are in accord with previous research (Beaver et al., 2016), and suggest that lower intelligence in general is associated with higher risk for victimization.

**Table 3.** Results from Binary Logistic Regression Models Predicting Victimization in the CNLSY

| Variable | Model 1 |  |  | Model 2 |  |  | Model 3 |  |  | Model 4 |  |  | Model 5 |  |  |
| --- | --- | --- | --- | --- | --- | --- | --- | --- | --- | --- | --- | --- | --- | --- | --- |
|  | OR | SE | 95% CI | OR | SE | 95% CI | OR | SE | 95% CI | OR | SE | 95% CI | OR | SE | 95% CI |
| **PIAT Reading Comprehension** |  |  |  |  |  |  |  |  |  |  |  |  |  |  |  |
| Standardized Measure | .97** | .01 | [.90-.99] | - | - | - | - | - | - | - | - | - | - | - | - |
| Bottom 25th Percentile | - | - | - | 1.10** | .02 | [1.01-1.68] | - | - | - | - | - | - | - | - | - |
| 50th Percentile | - | - | - | - | - | - | 1.02* | .03 | [1.00-1.92] | - | - | - | - | - | - |
| 75th Percentile | - | - | - | - | - | - | - | - | - | .94** | .01 | [.82-.98] | - | - | - |
| Top 25th Percentile | - | - | - | - | - | - | - | - | - | - | - | - | .89** | .01 | [.71-.95] |
| Family Income | .89** | .04 | [.75-.96] | .87* | .05 | [.71-.98] | .87* | .05 | [.70-.97] | .86* | .04 | [.74-.96] | .84* | .05 | [.71-.97] |
| Age | .98 | .02 | [.92-1.01] | .99 | .03 | [.89-1.01] | .99 | .03 | [.90-1.02] | .98 | .03 | [.87-1.02] | .98 | .03 | [.89-1.01] |
| Race | .92* | .02 | [.83-.99] | .93* | .05 | [.82-.99] | .92* | .05 | [.80-.99] | .92* | .05 | [.80-.99] | .92* | .05 | [.81-.99] |
| Sex | .95* | .01 | [.88-.99] | .96* | .01 | [.86-.99] | .95* | .01 | [.82-.99] | .95* | .01 | [.82-.99] | .95* | .01 | [.84-.99] |
| **PIAT Reading Recognition** |  |  |  |  |  |  |  |  |  |  |  |  |  |  |  |
| Standardized Measure | .99* | .03 | [.89-.99] | - | - | - | - | - | - | - | - | - | - | - | - |
| Bottom 25th Percentile | - | - | - | 1.06** | .02 | [1.01-1.72] | - | - | - | - | - | - | - | - | - |
| 50th Percentile | - | - | - | - | - | - | 1.01 | .03 | [.99-1.45] | - | - | - | - | - | - |
| 75th Percentile | - | - | - | - | - | - | - | - | - | .97* | .04 | [.69-.99] | - | - | - |
| Top 25th Percentile | - | - | - | - | - | - | - | - | - | - | - | - | .95** | .02 | [.78-.98] |
| Family Income | .91** | .03 | [.87-.96] | .89* | .03 | [.80-.97] | .93* | .02 | [.86-.98] | .98 | .02 | [.95-1.02] | .92* | .04 | [.84-.98] |
| Age | .99 | .02 | [.95-1.04] | .98 | .02 | [.96-1.03] | .98 | .01 | [.96-1.01] | .98 | .01 | [.97-1.02] | .97 | .02 | [.96-1.01] |
| Race | .93* | .05 | [.83-.98] | .94* | .03 | [.90-.99] | .98 | .01 | [.97-1.02] | .97 | .01 | [.96-1.01] | .96* | .03 | [.90-.99] |
| Sex | .96** | .01 | [.90-.98] | .97 | .02 | [.94-1.01] | .94* | .01 | [.89-.99] | .97 | .01 | [.93-1.03] | .95* | .02 | [.87-.99] |
| **PIAT Mathematics** |  |  |  |  |  |  |  |  |  |  |  |  |  |  |  |
| Standardized Measure | .97** | .02 | [.73-.99] | - | - | - | - | - | - | - | - | - | - | - | - |
| Bottom 25th Percentile | - | - | - | 1.15** | .01 | [1.02-1.52] | - | - | - | - | - | - | - | - | - |
| 50th Percentile | - | - | - | - | - | - | 1.04** | .02 | [1.01-1.38] | - | - | - | - | - | - |
| 75th Percentile | - | - | - | - | - | - | - | - | - | .90** | .02 | [.56-.98] | - | - | - |
| Top 25th Percentile | - | - | - | - | - | - | - | - | - | - | - | - | .79** | .01 | [.42-.95] |
| Family Income | .98 | .01 | [.90-1.02] | .96* | .03 | [.88-.98] | .95* | .01 | [.90-.99] | .98 | .02 | [.94-1.01] | .99 | .03 | [.95-1.06] |
| Age | .98 | .02 | [.96-1.03] | .99 | .02 | [.97-1.02] | .99 | .01 | [.97-1.05] | .97 | .01 | [.94-1.02] | .97 | .01 | [.94-1.02] |
| Race | .91** | .03 | [.84-.95] | .86** | .03 | [.78-.92] | .98 | .03 | [.96-1.02] | .95* | .03 | [.89-.98] | .88** | .02 | [.76-.96] |
| Sex | .97* | .01 | [.89-.99] | .98 | .02 | [.95-1.01] | .95* | .02 | [.88-.98] | .96* | .03 | [.87-.99] | .99 | .01 | [.96-1.03] |
| **PPVT-R** |  |  |  |  |  |  |  |  |  |  |  |  |  |  |  |
| Standardized Measure | .96** | .02 | [.90-.99] | - | - | - | - | - | - | - | - | - | - | - | - |
| Bottom 25th Percentile | - | - | - | 1.12** | .02 | [1.01-1.40] | - | - | - | - | - | - | - | - | - |
| 50th Percentile | - | - | - | - | - | - | 1.05* | .02 | [1.01-1.61] | - | - | - | - | - | - |
| 75th Percentile | - | - | - | - | - | - | - | - | - | .89** | .02 | [.72-.97] | - | - | - |
| Top 25th Percentile | - | - | - | - | - | - | - | - | - | - | - | - | .80** | .02 | [.69-.96] |
| Family Income | .97* | .01 | [.93-.99] | .92** | .02 | [.85-.97] | .95* | .01 | [.88-.99] | 1.01 | .01 | [.97-1.06] | .98 | .02 | [.95-1.03] |
| Age | .98 | .01 | [.95-1.02] | .98 | .01 | [.95-1.07] | .99 | .01 | [.97-1.01] | .99 | .01 | [.98-1.02] | .98 | .01 | [.96-1.01] |
| Race | .93** | .02 | [.84-.97] | .89* | .02 | [.80-.95] | .92** | .02 | [.83-.96] | .98 | .01 | [.96-1.02] | .99 | .02 | [.97-1.02] |
| Sex | .99 | .02 | [.97-1.04] | .98 | .03 | [.95-1.03] | .99 | .02 | [.97-1.02] | 1.00 | .01 | [.96-1.10] | .99 | .02 | [.94-1.06] |

Notes: CI = confidence interval. \*\* $p < .01$ ; \* $p < .05$

The next step in the analysis examines the genetic and environmental sources of variance that account for variance in different forms of intelligence and victimization in both sibling samples from the NLSY97 and CNLSY.^2^ Table 4 presents the between-sibling correlations for ASVAB scores and selfreported victimization for respondents from the NLSY97. Tetrachoric correlation coefficients across different types of sibling pairs show that MZ twin pairs displayed stronger concordance for ASVAB scores compared to DZ twin pairs, and that DZ twin pairs displayed stronger concordance for ASVAB scores compared to full siblings. Full siblings also displayed slightly stronger concordance for ASVAB scores compared to half-siblings. The same pattern of findings emerges for victimization among NLSY97 siblings. The between-sibling correlations, therefore, suggest that genetic influences account for some of the variance in liability for different quartiles of intelligence and victimization.

**Table 4.** Between-Sibling Correlations for Siblings from the NLSY97

|  | MZ Twins |  | DZ Twins |  | Full-Siblings |  | Half-Siblings |  |
| --- | --- | --- | --- | --- | --- | --- | --- | --- |
|  | *rho* | 95% *CI* | *rho* | 95% *CI* | *rho* | 95% *CI* | *rho* | 95% *CI* |
| <u>ASVAB</u> |  |  |  |  |  |  |  |  |
| Bottom 25th Percentile | .66 | [.34-.86] | .52 | [.29-.73] | .50 | [.27-.70] | .28 | [.09-.51] |
| 50th Percentile | .69 | [.42-.87] | .43 | [.21-.62] | .39 | [.21-.58] | .30 | [.12-.49] |
| 75th Percentile | .70 | [.40-.90] | .52 | [.27-.71] | .31 | [.17-.54] | .19 | [.05-.33] |
| Top 25th Percentile | .75 | [.58-.97] | .50 | [.20-.69] | .30 | [.15-.50] | .12 | [.02-.51] |
| <u>Victimization</u> | .81 | [.65-.93] | .72 | [.59-.82] | .38 | [.31-.48] | .20 | [.12-.29] |

Notes: *CI* = confidence interval

Table 5 presents the between-sibling tetrachoric correlation coefficients for PIAT scores, PPVT-R scores, and self-reported victimization for full- and half-siblings from the CNLSY. Estimates show that, with the exception of the bottom 25^th^ percentile of reading recognition, full-siblings display stronger concordance for reading comprehension, reading recognition, mathematics, and hearing and receptive vocabulary compared to half-siblings. Full-siblings also display stronger concordance for self-reported victimization compared to half-siblings. The between-sibling correlations, therefore, suggest that genetic influences account for some of the variance in liability for both intelligence and victimization. Taken together, the between-sibling correlations from the NLSY97 and CNLSY suggest that additive genetic influences may partially account for individual differences in liability for different thresholds of intelligence and risk for victimization.

**Table 5.** Between-Sibling Correlations for Siblings from the CNLSY

|  | Full-Siblings |  | Half-Siblings |  |
| --- | --- | --- | --- | --- |
|  | *rho* | 95% *CI* | *rho* | 95% *CI* |
| <u>PIAT Reading Comprehension</u> |  |  |  |  |
| Bottom 25th Percentile | .34 | [.21-.50] | .28 | [.18-.48] |
| 50th Percentile | .38 | [.24-.62] | .24 | [.13-.45] |
| 75th Percentile | .47 | [.30-.65] | .25 | [.12-.43] |
| Top 25th Percentile | .53 | [.23-.70] | .29 | [.17-.50] |
| <u>PIAT Reading Recognition</u> |  |  |  |  |
| Bottom 25th Percentile | .29 | [.11-.58] | .31 | [.15-.52] |
| 50th Percentile | .36 | [.18-.54] | .29 | [.19-.51] |
| 75th Percentile | .37 | [.19-.57] | .25 | [.11-.43] |
| Top 25th Percentile | .40 | [.26-.50] | .24 | [.09-.41] |
| <u>PIAT Mathematics</u> |  |  |  |  |
| Bottom 25th Percentile | .35 | [.17-.62] | .27 | [.16-.43] |
| 50th Percentile | .32 | [.20-.48] | .20 | [.08-.35] |
| 75th Percentile | .43 | [.26-.62] | .25 | [.10-.43] |
| Top 25th Percentile | .59 | [.22-.79] | .25 | [.08-.47] |
| <u>PPVT-R</u> |  |  |  |  |
| Bottom 25th Percentile | .38 | [.19-.53] | .25 | [.07-.42] |
| 50th Percentile | .34 | [.12-.47] | .20 | [.08-.34] |
| 75th Percentile | .41 | [.19-.57] | .18 | [.05-.37] |
| Top 25th Percentile | .42 | [.16-.59] | .17 | [.03-.31] |
| <u>Victimization</u> | .35 | [.23-.42] | .21 | [.14-.30] |

Notes: *CI* = confidence interval

We next estimate a series of univariate ACE models to examine the magnitude of additive genetic, shared environmental, and nonshared environmental effects on variance in liability for different percentiles of intelligence and risk for victimization. Table 6 presents the standardized parameter estimates from each estimated ACE model examining the genetic environmental effects on ASVAB scores and victimization among siblings from the NLSY97. Based on changes in χ^2^, CFI values, and RMSEA values, a CE model for ASVAB scores in the bottom 25^th^ percentile fit the data best, whereby shared environmental influences account for 36% of the variance in liability for below ASVAB scores in the bottom 25^th^ percentile, and nonshared environmental influences account for 64% of the variance in liability.^3^ Model fit indices indicate that AE models fit the data best when examining liability for ASVAB scores in the 50^th^ percentile, 75^th^ percentile, and top 25^th^ percentile of the distribution. Standardized parameter estimates indicate that additive genetic influences account for 34% of the variance in liability for scores in the 50^th^ percentile, 37% of the variance in liability for scores in the 75^th^ percentile, and 57% of the variance in liability for scores in the top 25^th^ percentile. Nonshared environmental influences therefore account for 66% of the variance in liability for scores in the 50^th^ percentile, 63% of the variance in liability for scores in the 75^th^ percentile, and 43% of the variance in liability for scores in the top 25^th^ percentile. For victimization, a full ACE model fit the data best, whereby additive genetic influences explain 24% of the variance in liability for victimization, shared environmental influences explain 16% of the variance in liability, and nonshared environmental influences explain the remaining 60% of the variance in liability.

**Table 6.** Parameter Estimates from Univariate Models Analyzing Sibling Pairs from the NLSY97

| | A | C | E | $\Delta\chi^2$ | $\Delta df$ | $p$ | CFI | RMSEA |
| --- | --- | --- | --- | --- | --- | --- | --- | --- |
| <u>ASVAB</u> |  |  |  |  |  |  |  |  |
| Bottom 25th Percentile |  |  |  |  |  |  |  |  |
| CE | .00 | .36*** | .64*** | 2.47 | 1 | .13 | .92 | .05 |
|  | [.00-.00] | [.16-.48] | [.52-.84] |  |  |  |  |  |
| 50th Percentile |  |  |  |  |  |  |  |  |
| AE | .34*** | .00 | .66*** | 1.78 | 1 | .21 | .93 | .05 |
|  | [.21-.56] | [.00-.00] | [.44-.79] |  |  |  |  |  |
| 75th Percentile |  |  |  |  |  |  |  |  |
| AE | .37*** | .00 | .63*** | 1.02 | 1 | .18 | .90 | .06 |
|  | [.20-.53] | [.00-.00] | [.47-.80] |  |  |  |  |  |
| Top 25th Percentile |  |  |  |  |  |  |  |  |
| AE | .57*** | .00 | .43*** | .75 | 1 | .27 | .89 | .06 |
|  | [.36-.70] | [.00-.00] | [.30-.64] |  |  |  |  |  |
| <u>Victimization</u> |  |  |  |  |  |  |  |  |
| ACE | .24** | .16* | .60*** | - | - | - | .94 | .04 |
|  | [.19-.32] | [.07-.23] | [.52-.71] |  |  |  |  |  |

Notes: Standardized ACE parameter estimates presented. 95% confidence intervals presented in brackets. CFI = comparative fit index, RMSEA = root mean square error of approximation. \*\*\* $p < .001$ ; \*\* $p < .01$ ; \* $p < .05$

Table 7 presents the standardized parameter estimates from each ACE model examining the genetic and environmental effects on PIAT scores from the CNLSY. An ACE model provided an adequate fit to the data for reading comprehension scores in the bottom 25^th^ percentile of the distribution. Based on this model, additive genetic influences explain 17% of the variance in liability for scores in the 25^th^ percentile, shared environmental influences explain 25% of the variance in liability, and nonshared environmental influences explain 58% of the variance in liability. An ACE model also fit the data best for reading comprehension scores in the 50^th^ percentile, whereby additive genetic influences explain 20% of the variance in liability, shared environmental influences explain 28% of the variance in liability, and nonshared environmental influences explain 52% of the variance in liability. With respect to reading comprehension scores in the 75^th^ percentile, an AE model was the best fitting model whereby additive genetic influences explain 41% of the variance in liability and nonshared environmental influences explain the remaining 59% of the variance in liability. A full ACE model fit the data best for reading comprehension scores in the top 25^th^ percentile, whereby additive genetic influences explain 19% of the variance in liability, shared environmental influence explain 25% of the variance in liability, and nonshared environmental influences explain 56% of the variance in liability.

ACE model results for reading recognition show that a CE model fit the data best when examining liability for reading recognition scores in the bottom 25^th^ percentile, while AE models fit the data best for reading recognition scores in the 50^th^ percentile, 75^th^ percentile, and top 25^th^ percentile. Results from the CE model indicate that shared environmental influences explain 27% of the variance in liability for reading recognition scores and nonshared environmental influences explain 73% of the variance in liability. Additive genetic influences explain 29% of the variance in liability for reading recognition scores in the 50^th^ percentile, 25% of the variance in liability for reading recognition scores in the 75^th^ percentile, and 46% of the variance in liability for reading recognition scores in the top 25^th^ percentile. Nonshared environmental influences are shown to explain 71% of the variance in liability for reading recognition scores in the 50^th^ percentile, 75% of the variance in liability for reading recognition scores in the 75^th^ percentile, and 54% of the variance in liability for reading recognition scores in the top 25^th^ percentile.

With respect to mathematics, model fit indices indicate that an AE model is the best fitting model for all quartiles of mathematical achievement. Based on the results from the best-fitting AE models, additive genetic influences explain 57% of the variance in liability for mathematic scores in the bottom 25^th^ percentile, 40% of the variance in liability for mathematic scores in the 50^th^ percentile, 64% of the variance in liability for mathematic scores in the 75^th^ percentile, and 70% of the variance in liability for mathematic scores in the top 25^th^ percentile. Nonshared environmental influences explain 43% of the variance in liability for mathematic scores in the bottom 25^th^ percentile, 60% of the variance in liability for mathematic scores in the 50^th^ percentile, 36% of the variance in liability for mathematics scores in the 75^th^ percentile, and 30% of the variance in liability for mathematic scores in the top 25^th^ percentile.

**Table 7.**
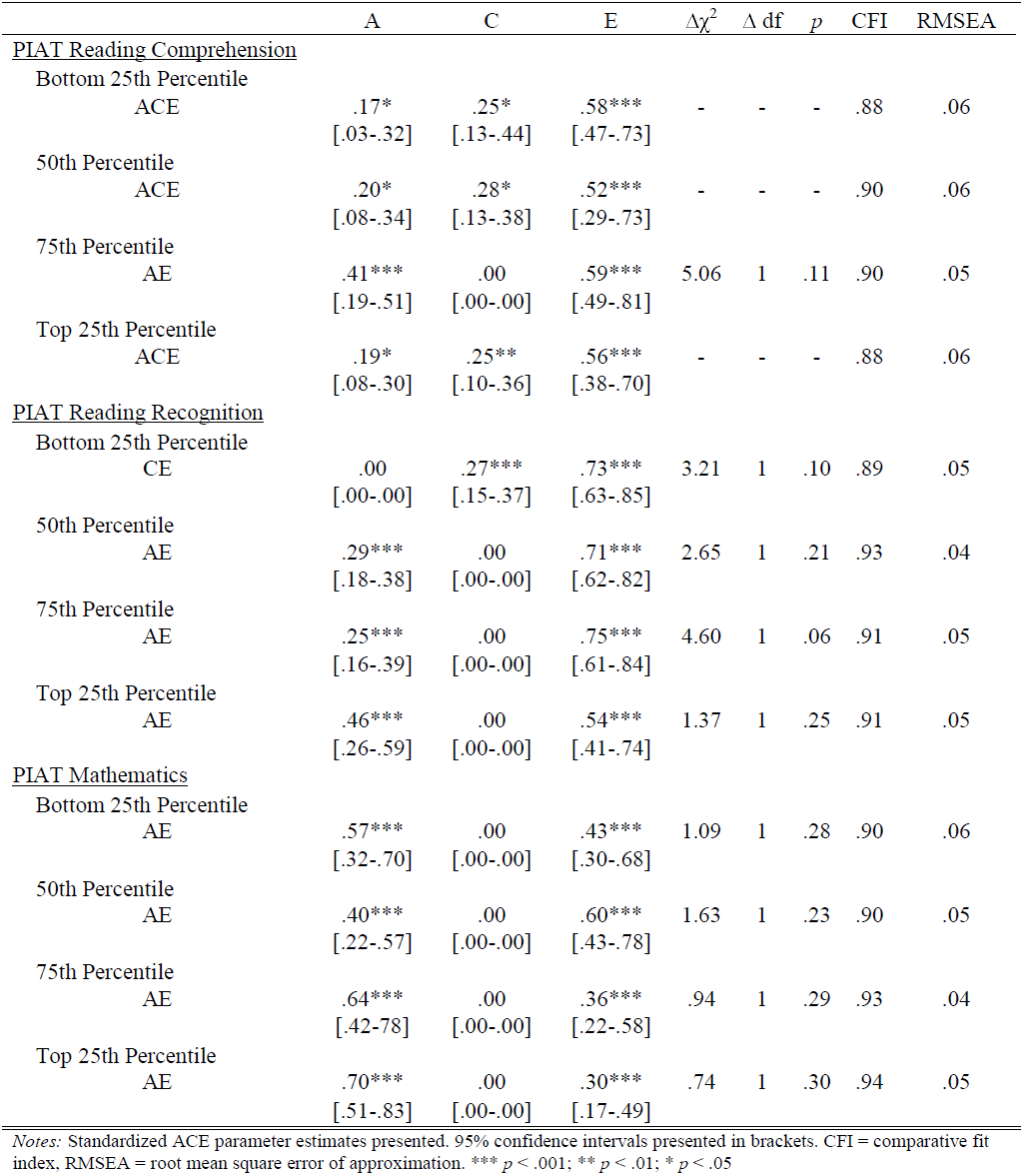
Parameter Estimates from Univariate Models Analyzing Sibling Pairs from the CNLSY

| | A | C | E | $\Delta\chi^2$ | $\Delta df$ | $p$ | CFI | RMSEA |
| --- | --- | --- | --- | --- | --- | --- | --- | --- |
| <u>PIAT Reading Comprehension</u> |  |  |  |  |  |  |  |  |
| Bottom 25th Percentile |  |  |  |  |  |  |  |  |
| ACE | .17*<br>[.03-.32] | .25*<br>[.13-.44] | .58***<br>[.47-.73] | - | - | - | .88 | .06 |
| 50th Percentile |  |  |  |  |  |  |  |  |
| ACE | .20*<br>[.08-.34] | .28*<br>[.13-.38] | .52***<br>[.29-.73] | - | - | - | .90 | .06 |
| 75th Percentile |  |  |  |  |  |  |  |  |
| AE | .41***<br>[.19-.51] | .00<br>[.00-.00] | .59***<br>[.49-.81] | 5.06 | 1 | .11 | .90 | .05 |
| Top 25th Percentile |  |  |  |  |  |  |  |  |
| ACE | .19*<br>[.08-.30] | .25**<br>[.10-.36] | .56***<br>[.38-.70] | - | - | - | .88 | .06 |
| <u>PIAT Reading Recognition</u> |  |  |  |  |  |  |  |  |
| Bottom 25th Percentile |  |  |  |  |  |  |  |  |
| CE | .00<br>[.00-.00] | .27***<br>[.15-.37] | .73***<br>[.63-.85] | 3.21 | 1 | .10 | .89 | .05 |
| 50th Percentile |  |  |  |  |  |  |  |  |
| AE | .29***<br>[.18-.38] | .00<br>[.00-.00] | .71***<br>[.62-.82] | 2.65 | 1 | .21 | .93 | .04 |
| 75th Percentile |  |  |  |  |  |  |  |  |
| AE | .25***<br>[.16-.39] | .00<br>[.00-.00] | .75***<br>[.61-.84] | 4.60 | 1 | .06 | .91 | .05 |
| Top 25th Percentile |  |  |  |  |  |  |  |  |
| AE | .46***<br>[.26-.59] | .00<br>[.00-.00] | .54***<br>[.41-.74] | 1.37 | 1 | .25 | .91 | .05 |
| <u>PIAT Mathematics</u> |  |  |  |  |  |  |  |  |
| Bottom 25th Percentile |  |  |  |  |  |  |  |  |
| AE | .57***<br>[.32-.70] | .00<br>[.00-.00] | .43***<br>[.30-.68] | 1.09 | 1 | .28 | .90 | .06 |
| 50th Percentile |  |  |  |  |  |  |  |  |
| AE | .40***<br>[.22-.57] | .00<br>[.00-.00] | .60***<br>[.43-.78] | 1.63 | 1 | .23 | .90 | .05 |
| 75th Percentile |  |  |  |  |  |  |  |  |
| AE | .64***<br>[.42-.78] | .00<br>[.00-.00] | .36***<br>[.22-.58] | .94 | 1 | .29 | .93 | .04 |
| Top 25th Percentile |  |  |  |  |  |  |  |  |
| AE | .70***<br>[.51-.83] | .00<br>[.00-.00] | .30***<br>[.17-.49] | .74 | 1 | .30 | .94 | .05 |

Notes: Standardized ACE parameter estimates presented. 95% confidence intervals presented in brackets. CFI = comparative fit index, RMSEA = root mean square error of approximation. \*\*\* $p < .001$ ; \*\* $p < .01$ ; \* $p < .05$

Table 8 presents the standardized parameter estimates from each univariate ACE model assessing the magnitude of genetic and environmental effects on PPVT-R scores and victimization. An ACE model fit the data best for PPVT-R scores in the bottom 25^th^ percentile of distribution whereby additive genetic influences explain 47% of the variance in liability for PPVT-R scores in the bottom 25^th^ percentile, shared environmental influences explain 19% of the variance in liability, and nonshared environmental influences explain 34% of the variance in liability. Additive genetic and nonshared environmental influences best explained variance in liability for PPVT-R scores in the 50^th^, 75^th^, and top 25^th^ percentile. Additive genetic influences explain 41% of the variance in liability for PPVT-R scores in the 50^th^ percentile, 45% of the variance in liability for PPVT-R scores in the 75^th^ percentile, and 61% of the variance in liability for PPVT-R scores in the top 25^th^ percentile. For victimization, the AE model fit the data best, whereby additive genetic influences explain 37% of the variance in liability for being the victim of a violent crime, and nonshared environmental factors explain the remaining 63% of the variance in liability.

**Table 8.** Parameter Estimates from Univariate Models Analyzing PPVT-R Scores and Victimization in the CNLSY

| | A | C | E | $\Delta\chi^2$ | $\Delta df$ | $p$ | CFI | RMSEA |
| --- | --- | --- | --- | --- | --- | --- | --- | --- |
| <u>PPVT-R</u> |  |  |  |  |  |  |  |  |
| Bottom 25th Percentile |  |  |  |  |  |  |  |  |
| ACE | .47**<br>[.28-.63] | .19*<br>[.10-.32] | .34***<br>[.21-.49] | - | - | - | .91 | .05 |
| 50th Percentile |  |  |  |  |  |  |  |  |
| AE | .41***<br>[.26-.59] | .00<br>[.00-.00] | .59***<br>[.41-.74] | 1.63 | 1 | .17 | .93 | .03 |
| 75th Percentile |  |  |  |  |  |  |  |  |
| AE | .45***<br>[.32-.58] | .00<br>[.00-.00] | .55***<br>[.42-.68] | 3.80 | 1 | .06 | .92 | .05 |
| Top 25th Percentile |  |  |  |  |  |  |  |  |
| AE | .61***<br>[.45-.80] | .00<br>[.00-.00] | .39***<br>[.20-.55] | 1.93 | 1 | .15 | .93 | .04 |
| <u>Victimization</u> |  |  |  |  |  |  |  |  |
| AE | .37***<br>[.28-.49] | .00<br>[.00-.00] | .63***<br>[.54-.72] | .07 | 1 | .62 | .93 | .06 |

Notes: Standardized ACE parameter estimates presented. 95% confidence intervals presented in brackets. CFI = comparative fit index, RMSEA = root mean square error of approximation. \*\*\* $p < .001$ ; \*\* $p < .01$ ; \* $p < .05$

The next step of the analysis involved estimating the extent to which genetic and environmental sources account for the covariance between different quartiles of intelligence and victimization in both the NLSY97 and CNLSY sibling samples. Table 9 presents the standardized parameter estimates from models assessing the genetic and environmental overlap between different quartiles of ASVAB scores and victimization.^4^ After comparing the baseline bivariate ACE model to other nested models for the association between ASVAB scores in the bottom 25^th^ percentile and victimization, model fit indices indicate that the CE model fit the data best. Parameter estimates from the best-fitting CE bivariate liability-threshold model reveal that 43% of the covariance in liability between ASVAB scores in the bottom 25^th^ percentile and risk for victimization is accounted for by shared environmental influences, whereas 57% of the covariance in liability is accounted for by nonshared environmental influences. AE models fit the data best for all other bivariate models, showing that additive genetic influences explain 34% of the covariance in liability between ASVAB scores in the 50^th^ percentile and victimization, 30% of the covariance in liability between ASVAB scores in the 75^th^ percentile and victimization, and 58% of the covariance in liability between ASVAB scores in the top 25^th^ percentile and victimization. Nonshared environmental influences therefore explain 66% of the covariance in liability between ASVAB scores in the 50^th^ percentile and victimization, 70% of the covariance in liability between ASVAB scores in the 75^th^ percentile and victimization, and 42% of the covariance in liability between ASVAB scores in the top 25^th^ percentile and victimization.

**Table 9.**
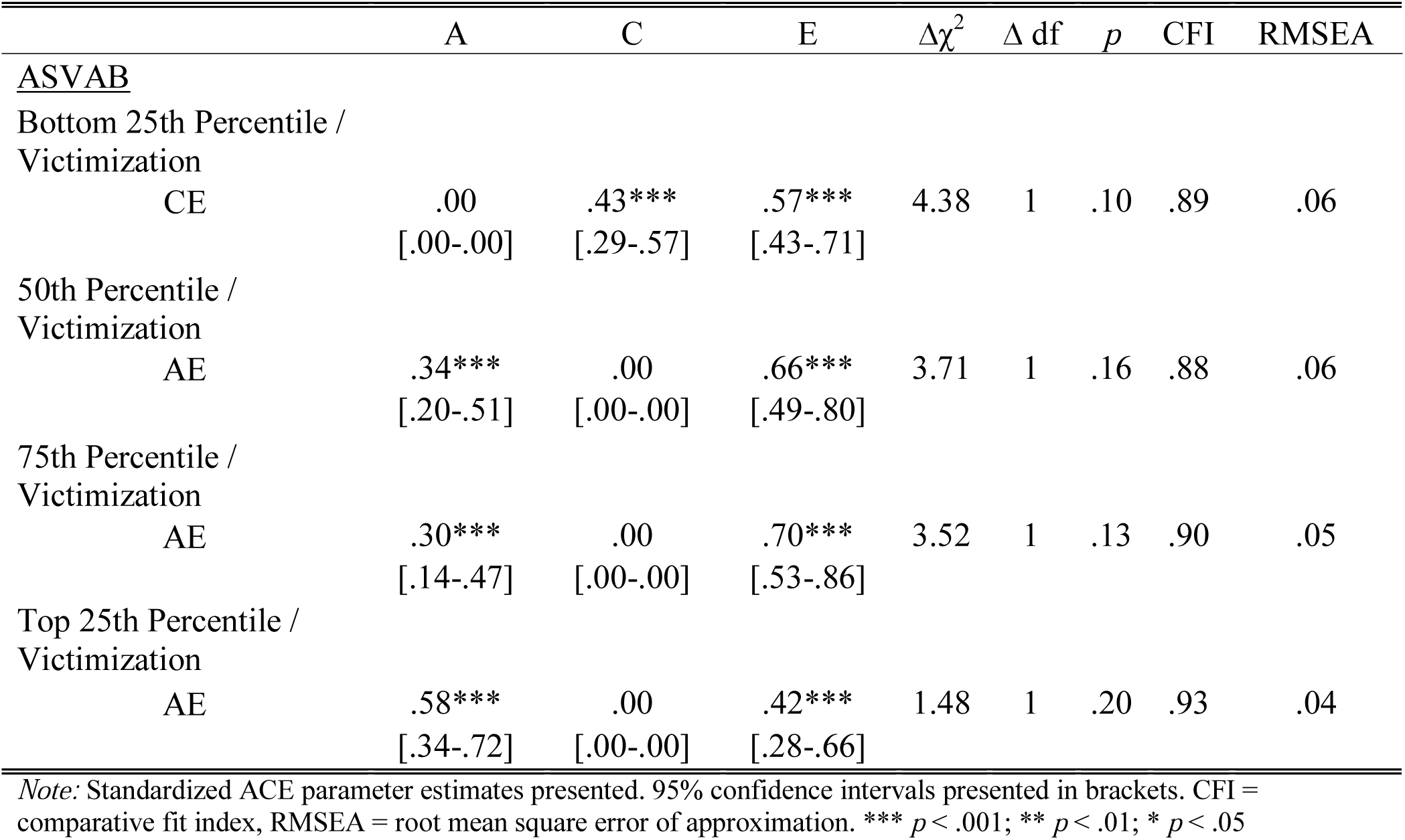
Parameter Estimates from Bivariate Liability-Threshold Models for ASVAB Scores and Victimization

Table 10 presents the standardized parameter estimates from each bivariate liability-threshold model examining the genetic and environmental effects on the covariance between PIAT scores and victimization. Based on model fit indices, an AE model was the best-fitting model for the association between reading comprehension scores in the 25^th^ percentile, 75^th^ percentile, top 25^th^ percentile, and victimization. A full ACE model fit the data best for reading comprehension scores in the 50^th^ percentile and victimization. Additive genetic influences were found to explain 27% of the covariance in liability between reading comprehension scores in the 25^th^ percentile and victimization, 12% of the covariance in liability between reading comprehension scores in the 50^th^ percentile and victimization, 58% of the covariance in liability between reading comprehension scores in the 75^th^ percentile and victimization, and 34% of the covariance in liability between reading comprehension scores in the top 25^th^ percentile and victimization. Shared environmental influences were found to only explain 15% of the covariance in liability between reading comprehension scores in the 50^th^ percentile and victimization. Model fit indices suggest that a CE model fit the data best for the association between scores in the bottom 25^th^ percentile of the reading recognition measure and victimization, while AE models are the best-fitting models for the association between scores in the 50^th^, 75^th^, and top 25^th^ percentile, and victimization. Shared environmental influences explain 37% of the covariance in liability between reading recognition scores in the bottom 25^th^ percentile and victimization, whereas nonshared environmental influences explain 63% of the covariance in liability between reading recognition scores in the bottom 25^th^ percentile and victimization.

Additive genetic influences explained 41% of the covariance in liability between reading recognition scores in the 50^th^ percentile and risk for victimization, 62% of the covariance in liability between reading recognition in the 75^th^ percentile and risk for victimization, and 67% of the covariance in liability between reading recognition in the top 25^th^ percentile and risk for victimization. Bivariate analyses examining the covariance between liability for different quartiles of mathematics and risk for victimization show that AE models fit the data best. Parameter estimates from the best-fitting bivariate models show that additive genetic influences explain 52% of the covariance in liability between mathematics scores in the bottom 25^th^ percentile and risk for victimization, 24% of the covariance in liability between mathematics scores in the 50^th^ percentile and risk for victimization, 65% of the covariance in liability between mathematics scores in the 75^th^ percentile and risk for victimization, and 69% of the covariance in liability between mathematics scores in the top 25^th^ percentile and risk for victimization.

**Table 10.**
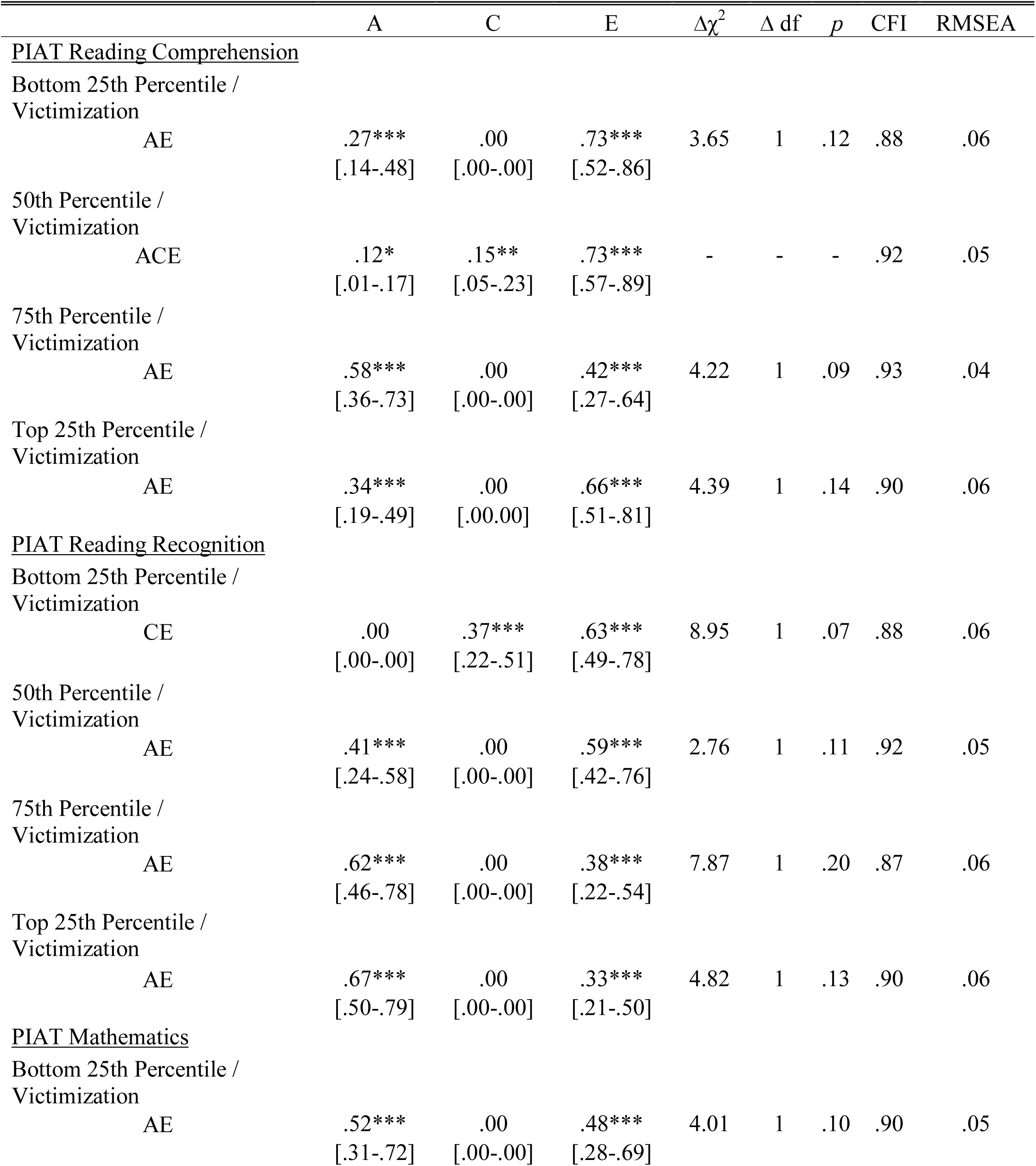
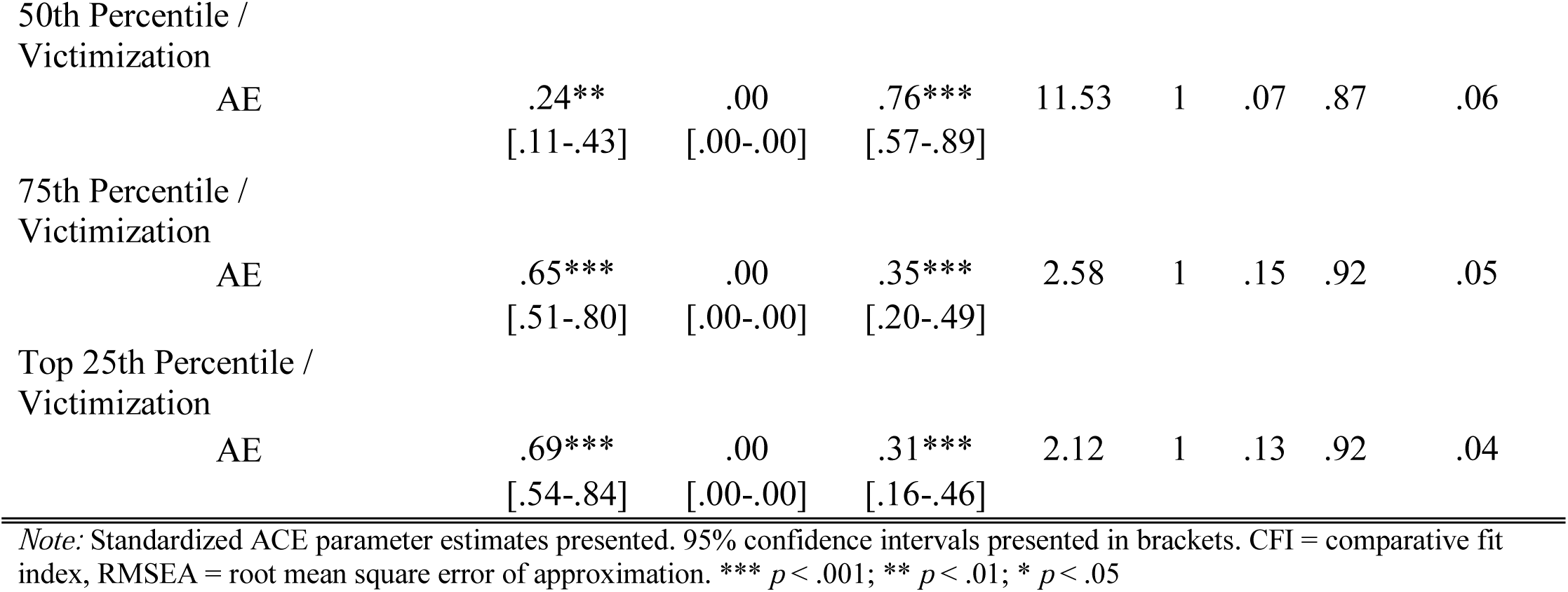
Parameter Estimates from Bivariate Liability-Threshold Models for PIAT Reading Scores and Victimization

Table 11 presents the final set of bivariate liability-threshold models examining the genetic and environmental overlap between different quartiles of PPVT-R scores and victimization. Model fit indices suggest that AE models fit the data best. The results from the best-fitting AE models indicate that additive genetic influences explain 20% of the covariance in liability between hearing and receptive vocabulary in the bottom 25^th^ percentile and risk for victimization, 18% of the covariance in liability between hearing and receptive vocabulary in the 50^th^ percentile and risk for victimization, 44% of the covariance in liability between hearing and receptive vocabulary in the bottom 75^th^ percentile and risk for victimization, and 57% of the covariance in liability between hearing and receptive vocabulary in the top 25^th^ percentile and risk for victimization. Nonshared environmental influences explain the remaining covariance between each quartile of hearing and receptive vocabulary and victimization.

**Table 11.** Parameter Estimates from Bivariate Liability-Threshold Models for PPVT-R Scores and Victimization

| | A | C | E | $\Delta\chi^2$ | $\Delta df$ | $p$ | CFI | RMSEA |
| --- | --- | --- | --- | --- | --- | --- | --- | --- |
| <u>PPVT-R</u> |  |  |  |  |  |  |  |  |
| Bottom 25th Percentile / Victimization |  |  |  |  |  |  |  |  |
| AE | .20***<br>[.07-.38] | .00<br>[.00-.00] | .80***<br>[.62-.93] | 10.48 | 1 | .07 | .88 | .06 |
| 50th Percentile / Victimization |  |  |  |  |  |  |  |  |
| AE | .18***<br>[.08-.32] | .00<br>[.00-.00] | .82***<br>[.68-.92] | 8.54 | 1 | .11 | .89 | .06 |
| 75th Percentile / Victimization |  |  |  |  |  |  |  |  |
| AE | .44***<br>[.21-.59] | .00<br>[.00-.00] | .56***<br>[.41-.79] | 7.32 | 1 | .10 | .89 | .06 |
| Top 25th Percentile / Victimization |  |  |  |  |  |  |  |  |
| AE | .57***<br>[.38-.79] | .00<br>[.00-.00] | .43***<br>[.21-.62] | 2.03 | 1 | .19 | .92 | .04 |

*Note:* Standardized ACE parameter estimates presented. 95% confidence intervals presented in brackets. CFI = comparative fit index, RMSEA = root mean square error of approximation. \*\*\* $p < .001$ ; \*\* $p < .01$ ; \* $p < .05$

The last step of the analysis involved estimating a series of within-sibling regressions to examine the effect of intelligence on victimization above and beyond the influence of within-family confounds such as additive genetic and shared environmental factors. Table 12 reports the results from the estimated regression models. The results reveal that a twin or sibling with a higher intelligence score is less likely to report being the victim of violent crime as compared to their co-twin or co-sibling with a lower intelligence score. The effect of intelligence on victimization was significant and comparable in size across sibling category in both samples, suggesting that intelligence exerts a consistent influence on risk for victimization after controlling for a range of relevant within family confounds.

**Table 12.** Within-Sibling Regression Analysis Predicting Victimization

|  | MZ Twins | DZ Twins | Full-Siblings | Half-Siblings |
| --- | --- | --- | --- | --- |
| <u>NLSY97</u> |  |  |  |  |
| ASVAB | .91* | .93* | .87*** | .90*** |
|  | [.52-.99] | [.58-.98] | [.69-.96] | [.81-.97] |
| <u>CNLSY</u> |  |  |  |  |
| PIAT Reading Comprehension | - | - | .92*** | .93** |
|  |  |  | [.79-.98] | [.76-.99] |
| PIAT Reading Recognition | - | - | .90*** | .93** |
|  |  |  | [.75-.97] | [.71-.99] |
| PIAT Mathematics | - | - | .96*** | .98* |
|  |  |  | [.78-.98] | [.69-.99] |
| PPVT-R | - | - | .91*** | .96* |
|  |  |  | [.78-.97] | [.75-.99] |

Notes: Presented estimates are odds ratios. 95% confidence intervals in brackets.
\*\*\* $p <$
.001; \*\* $p < .01$ ; \* $p < .05$

## Discussion

Research investigating the correlates of victimization has identified a number of individual-level factors influence the likelihood of victimization, including indicators of general intelligence (Beaver et al., 2016). The current study sought to replicate and extend Beaver and colleague’s findings using two nationally representative samples of American youth. In addition to examining the covariance between intelligence and victimization, the use of a genetically sensitive research designs allowed us to examine the extent to which the phenotypic association between intelligence and victimization is accounted for by a shared genetic etiology between the two traits.

Our findings revealed an association between lower intelligence (on average) and victimization in both the National Longitudinal Survey of Youth 1997 (NLSY97) and the Children of the National Longitudinal Survey of Youth 1979 (CNLSY). It is worth noting, that in terms of effect size the magnitude of the intelligence-victimization association in both samples, while consistent, was not large.^5^ In general, however, our results accord with those of Beaver et al. (2016) who also documented an association between intelligence and victimization. Our findings also extend prior research in this area in several important respects. First, the current study used broader measures of intelligence in both the NLSY97 sample—which assessed arithmetic reasoning, mathematics, paragraph comprehension, and word knowledge—and the CNLSY sample—which assessed reading comprehension and recognition, mathematics, and vocabulary. Given that our results show robust associations between victimization and different measures of intelligence, in combination with previous research reporting an association between victimization and verbal intelligence (Beaver et al., 2016), we can have greater confidence in the intelligence-victimization link.

The findings of the current study, moreover, elucidate the extent to which covariance between different types of intelligence and criminal victimization is accounted for by shared genetic and environmental factors underpinning both traits. The general pattern of our results suggests that a combination of genetic and environmental factors, common to both intelligence and victimization, best explained the association between the traits. The fact that intelligence and victimization may share a genetic etiology, and that both traits are heritable (Beaver et al. 2009; Plomin & Deary, 2015), however, points to the importance of accounting for genetic influences on variables of interest in subsequent research. The influence of intelligence on victimization remained statistically significant in the models, after correcting for the possibility of familial confounding. Future research in this area, then, should ensure properly accounting for genetic factors when examining the influence of general intelligence on victimization experiences (see Barnes, Boutwell, Beaver, Gibson & Wright, 2014).

It is important to note some of the limitations of our study. First, the use of twin and sibling samples may limit the generalizability of the current findings to populations of non-twin/non-sibling participants. Barnes and Boutwell (2013), however, provided reason to suspect that twins and non-twins/non-siblings are not substantively different from one another across a host of phenotypes. Second, some of the analysis included intelligence measures that divided respondents into percentile groups based on their scores across the various measure of intelligence. Percentile scores afforded the assessment of the effect of intelligence separately across different levels of ability, therefore allowing the observation of how intelligence impacts the odds of victimization (either positively in the lower percentiles, or negatively in the higher percentiles). Nonetheless, there are tradeoffs to categorizing a continuous measure (e.g., losing natural variation in the measure). Although we retained the continuous measure for the within-sibling regression analyses, we were unable to do so for the ACE model analyses due to the dichotomous nature of the victimization item.^6^ Future research with access to continuous measures of victimization should attempt to replicate our findings. Finally, our measure of victimization captured only the general experience of victimization, and was therefore incapable of distinguishing participants who may have experienced repeat victimization. Future studies utilizing alternative measures of victimization may reveal that the current findings vary as the frequency of victimization increases—this, however, remains an open empirical question.

The current research provides further evidence for the association between lower intelligence and increased risk of personal victimization that accord with previous research on the topic (Beaver et al., 2016). We also extended this association to cover a wider array of intelligence assessments using independent samples from prior studies. Our results from two national samples of Americans suggest that the association between intelligence and victimization can be explained, in part, by a shared genetic etiology underpinning both traits. These findings represent an important step toward the integration of behavioral genetic analyses into criminological and psychological theories of victimization, and further highlight the importance of accounting for genetic influences when studying victimization experiences (Barnes et al., 2014; McGue et al., 2010).

## Footnotes

1 All bivariate associations between indicators of intelligence and victimization were statistically significant (even if they did not reach significance in the multivariate model). As a result, ACE models were estimated across all percentile levels for intelligence variables.

2 Cross-twin cross-trait correlations are available upon request.

3 In order to preserve space, we present only the best fitting models here. However, interested readers can consult supplementary information for complete tables, containing all possible model iterations.

4 Full ACE models were not estimated in the bivariate analysis when the shared environmental parameter was non-significant in the best fitting univariate ACE models for the measures of intelligence and victimization.

5 An additional point worth making is that individuals scoring in the lowest 25^th^ percentile on the ASVAB measures were *not* more likely to be victimized. A finding that may deserve more investigation in future research.

6 Some sibling based modeling strategies will accommodate a continuous and dichotomous variable. However, these approaches can present with certain challenges—including unstable parameter estimates and poor model fit. With this in mind, we opted to retain our approach in the current analysis.

